# Brain organoid computing for robotic decision-making

**DOI:** 10.64898/2026.09.09.750426

**Authors:** Hongwei Cai, Chunhui Tian, Yang Yang, Yantao Xing, Zichen Hong, Huiyu Chu, Jiansen Wang, Zheng Ao, Jason S. Meyer, James Friend, Jason Tchieu, Mingxia Gu, Insoo Hyun, Ken Mackie, Lantao Liu, Feng Guo

## Abstract

Biomimicry has inspired the evolution of robotics toward greater autonomy, adaptability, and symbiosis with humans and dynamic environments. However, current robotic systems still face major challenges in recapitulating the high-efficiency decision-making capabilities of the human brain under complex and dynamic conditions. Here, we present Brainobot, a biohybrid robotic system that establishes a brain organoid controller as a high-level robotic decision-making layer for closed-loop embodiment. By leveraging brain organoid reservoir computing, Brainobot interacts with dynamic environments by receiving and processing sensory inputs and generating motor actions. As a proof-of-concept demonstration, Brainobot is implemented in a humanoid robotic system to perform real-world tasks, including object grasping and laser chasing. Interestingly, Brainobot exhibits unique features, including cross-task adaptivity, high computing efficiency, and low energy consumption. Thus, our approach may provide insights for advancing robotic embodiment and understanding biological decision-making.

---

The evolution of robotics has been dramatically changing modern society from industry and healthcare to environmental monitoring, space exploration, transportation, and manufacturing(*1–5*). This transformation is mainly driven by advances in robotic autonomy and system embodiments that enable robots to sense, process, and act with complex, dynamic environments and humans(*6–8*). Classical decision-making frameworks have been dominated by the perception–planning–control pipeline which has achieved substantial progress over the past two decades, particularly in high-precision, time-critical tasks within structured environments. (*9–11*). However, significant challenges remain to use classical decision-making theories to faithfully address real-world complexity and dynamics. By leveraging AI models including deep learning and reinforcement learning, adaptive and data-driven strategies recently have been well adopted for robotic decision making, enabling human-like performance in handling challenging real-world tasks(*1, 7, 12–15*). Despite these promising advances, such systems are still constrained by high energy consumption, data-intensive training, and limited cross-task adaptivity to interact with unstructured environments(*6, 16–20*). Thus, significant gaps remain, necessitating advances in control paradigms to support efficient robotic decision-making in unstructured, real-world environments.

Biomimicry may provide unique opportunity and solution, since the human brain represents a fundamentally different computational paradigm that avoids the von Neumann bottleneck limiting current silicon-based hardware systems(*21–23*). It leverages functional biological neural networks to continuously process complex spatiotemporal information, enabling rapid, context-aware decision-making with remarkable energy and data efficiency(*24–26*). Moreover, such systems exhibit intrinsic advantages, including adaptability and resilience(*27*), arising from the dynamics of biological neural networks(*28–30*). Inspired by these principles, bio-robotics has begun to incorporate biological components, such as two-dimensional cultures of dissociated neurons, into robotic decision-making frameworks(*31–33*). Moving beyond two-dimensional culture systems, human brain organoids, three-dimensional brain-like tissues derived from human stem cells, further advance this paradigm by recapitulating key cellular, structural, and functional features of complex human neural networks(*34–39*). Building on this, biohybrid computing systems leverage the computational capacity of organoid neural networks to emulate aspects of human high-efficiency information processing(*40–43*). Our recent work, Brainoware, has provided a demonstration of brain organoid reservoir computing, enabling tasks such as speech recognition and nonlinear prediction(*40*). Recently, complementary studies have advanced the maturation, interfacing, and functional assessment of brain organoids toward directing robotic movements(*44*) and have demonstrated goal-directed learning through feedback-driven neural plasticity(*45*). Despite these advances, the application of organoid computing for real-world robotic decision-making remains largely unexplored.

Here, we present Brainobot, a biohybrid robotic system that integrates an organoid controller as a high-level robotic decision-making layer. We implement the system into a humanoid to establish a closed-loop sense–compute–action system. We demonstrate that Brainobot can process real-time sensory inputs and generate motor outputs to perform real-world tasks in dynamic environments such as object grasping and laser chasing, showing unique features such as cross-task adaptivity, data efficiency, and energy efficiency.

## Results

### Brainobot that leverages organoid computing for robotic decision-making

We propose Brainobot, a biohybrid robotic system that uses a brain organoid processor for high-level robotic decision-making. As a proof-of-concept demonstration, we implemented Brainobot to control a humanoid robot by leveraging reservoir computing within the organoid processor (**Fig.1a**). In this humanoid system, the organoid processor receives dynamic environmental information, computes high-level commands, and transmits these commands to mid-/low-level controllers to drive corresponding motor actions. This configuration establishes bidirectional information exchange between the biological and robotic domains, enabling a closed-loop sense-compute-action framework for robotic decision-making across various real-world tasks (**Supplementary Fig.1**). The hypothesized learning curve of Brainobot illustrates progressive improvements in task performance and cross-task adaptivity over time, potentially driven by experience-dependent reshaping of organoid neural networks through neural plasticity (**Fig.1b**). In our experiments, the organoid processor was employed for remote control of a humanoid (**Fig.1c**), establishing a prototype platform for embodied interaction in real-world environments. The organoid processor was constructed by mounting a functional cortical organoid onto a MaxOne microelectrode array (MEA) chip (**Fig.1d**), enabling bidirectional information exchange with the humanoid system. Within the organoid processor, functional neural networks in the cortical organoid are essential for these bidirectional information exchange. To ensure the development of functional organoid neural networks, we characterized the maturation of cortical organoids (**Supplementary Fig.2**). We found that 3-month-old cortical organoids (**Fig.1e**) exhibited densely interconnected neural networks and formed complex three-dimensional neural architectures containing mature neurons (MAP2^+^) and astrocytes (GFAP^+^) (**Fig.1f**). Together, these results establish the structural and functional foundation to develop organoid processors for high-level robotic decision-making in humanoid systems. Then, we integrated and validated Brainobot to conduct various tasks and to demonstrate its unique features in interacting with real-world environments.

**Figure 1.**
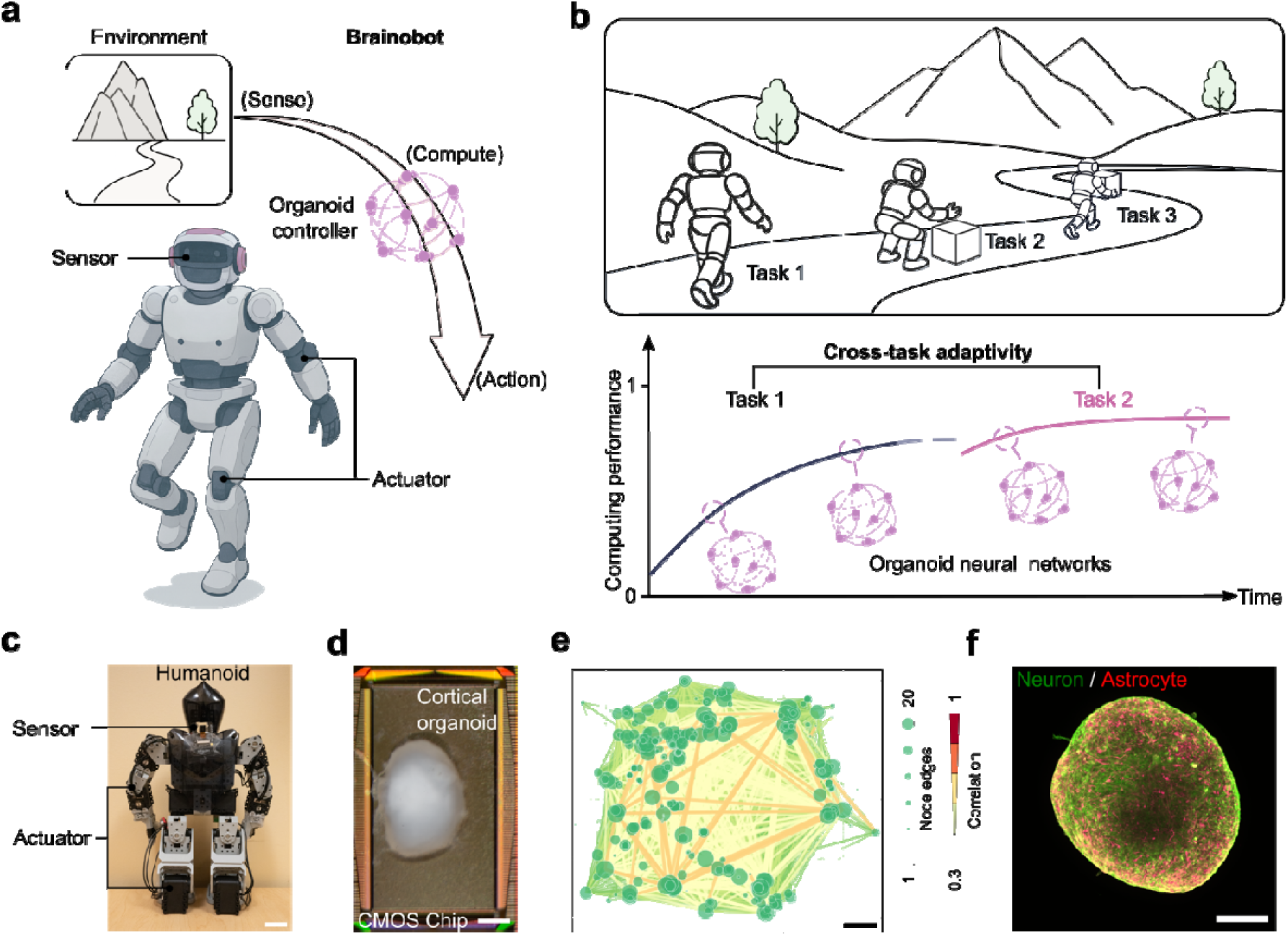
Brainobot that leverages organoid computing for robotic decision-making. **a.** Schematic diagram of the Brainobot framework for controlling a humanoid. The organoid controller receives sensory inputs from the humanoid, processes the information, and outputs action commands that drive corresponding movements through organoid neural networks. **b.** Schematic illustration of Brainobot performing real-world tasks and the hypothesized corresponding learning curves over time and across different tasks through reshaping of organoid neural networks. **c.** Image of a humanoid implemented with remote control of an organoid controller. **d.** Physical setup of an organoid controller by mounting a cortical organoid onto a MaxOne microelectrode array (MEA) chip, scale bar = 500μm. **e.** Representative functional connectivity map of a 3-month-old cortical organoid on MEA, scale bar = 200μm. Edge color and width indicate the degree of correlation between neuron units, while node size reflects the number of connections per neuron. **f.** Whole-mount immunostaining of a 3-month-old cortical organoid showing complex three-dimensional neuronal networks with mature neurons (MAP2_+_) and astrocytes (GFAP_+_), scale bar = 200pm.

### System integration and validation

As a proof-of-concept application in a humanoid system, we integrated the hardware and software components of Brainobot and validated the system (**Supplementary Fig.3**) using a simple task. The hardware system (**Fig.2a**) was established by implementing Brainobot on a commercially available humanoid robot platform (ROBOTIS Premium), which is equipped with a camera and a distance sensor for capturing environmental information, servo motors for actuation, and an onboard controller for mid- and low-level motor execution. In this architecture, the organoid processor received processed sensory features from the humanoid and generated output signals corresponding to predefined motor actions (e.g., hand raising) and locomotion behaviors (e.g., walking). The onboard mid/low-level controller then translated these high-level output commands into low-level control signals for individual servo motors, enabling physical execution of the motor actions. The software system (**Fig.2b**) operated under a reservoir computing framework based on the organoid processor. Environmental inputs were transformed into spatiotemporal electrical stimulation sequences through an input encoding layer and delivered to the brain organoid via the MEA interface. These inputs were projected into a high-dimensional computational space through the nonlinear dynamics and fading-memory properties of the organoid neural networks. The resulting evoked neural activity was recorded and decoded through a readout function to generate high-level motor commands that drove the low-level control layer of the humanoid.

**Figure 2.**
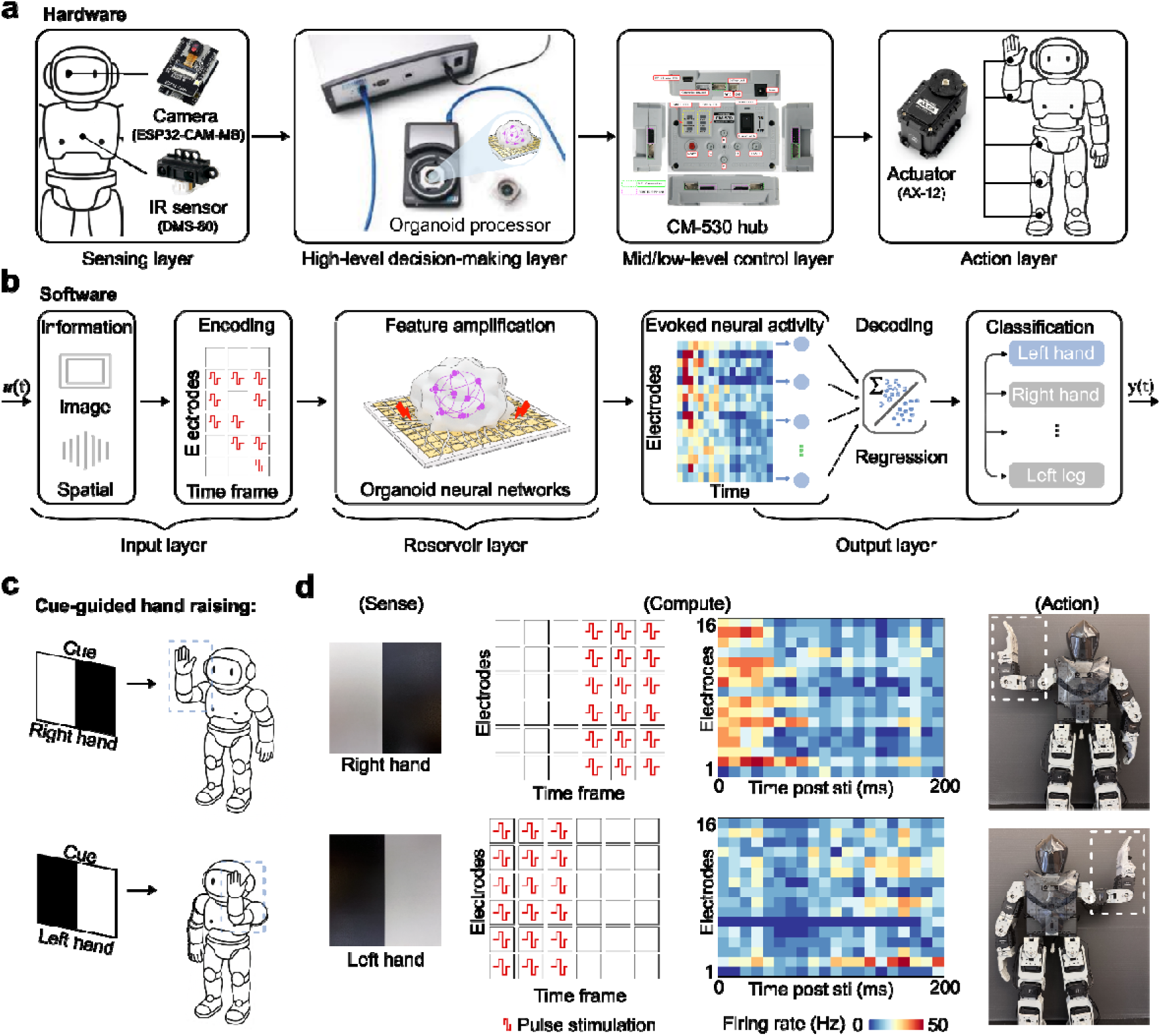
System integration and validation of Brainobot. **a.** Hardware architecture of Brainobot. **b.** Software framework of Brainobot. **c.** Design of the cue-guided hand-raising task. **d.** Conduction of the cue-guided hand-raising task through a sense-compute-action framework. Representative images of visual cues captured by the camera of the humanoid. Temporospatially encoded stimulation patterns corresponding to these two visual cues, together with representative raster plots showing the evoked neural activity in response to the corresponding stimulation patterns. Representative images of the humanoid raising the right/left hand in response to the corresponding visual cues.

After system integration, we validated the prototype using a simple cue-guided hand-raising task (**Fig.2c**). The humanoid was programmed to raise its right hand when presented with a visual cue (left: white; right: black), and to raise its left hand when presented with the reversed visual cue (left: black; right: white). We further evaluated the system using this task by tracking and optimizing each step of its sense-compute-action loop (**Fig.2d, Supplementary Figs.4 and 5**). Visual input (400 × 300 pixels) captured by the onboard camera was downsampled and brightness-thresholded into a 6 × 6 binary matrix, which was subsequently converted into a bipolar stimulation pulse sequence (6 electrodes × 6 time steps) and delivered to the organoid through the MEA interface. The evoked neural responses (e.g., raster plots) were recorded and used as input features for a logistic regression classifier. After training the logistic regression model and optimizing the stimulation patterns, the organoid processor successfully performed the hand-raising task in response to the appropriate visual cues, demonstrating its ability to interpret environmental inputs and generate task-specific motor outputs. Then, we demonstrated advanced tasks of this system.

### Grasping task

We further evaluated the capability of Brainobot for decision-making under more complex sensory inputs involving multiple classes. Specifically, we designed an object-specific grasping task that required selective motor responses to distinct visual cues (**Fig.3a**). In this task, the humanoid was programmed to grasp the letter block “I” using its left hand, grasp the letter block “U” using its right hand, or remain idle when no block was presented. We evaluated the system performance in this task by tracking and optimizing each step of the sense-compute-action loop (**Fig.3b, Supplementary Fig.6, Supplementary Video.1**). Visual images captured by the humanoid’s onboard camera were preprocessed into binary feature maps representing object identity and location. These sensory features were then transformed into temporospatially encoded stimulation patterns and delivered to the cortical organoid in the organoid processor through the MEA interface. Distinct patterns of evoked neural activity were observed in response to the three stimulation sequences, as shown in the representative raster plots, indicating classification of the visual inputs by the organoid biocontroller based on its internal network dynamics. The decoded neural outputs were subsequently mapped to corresponding motor commands, driving appropriate grasping behaviors in the humanoid robot. To evaluate the functional contribution of the organoid neural networks, we compared system performance under normal conditions and under pharmacological inhibition such as with tetrodotoxin (TTX) treatment (**Fig.3c**). The confusion matrix revealed that the organoid processor achieved high accuracy in associating sensory cues with the correct motor actions, whereas TTX treatment, which blocks neuronal activity, drastically impaired decision-making performance. Quantitative analysis (**Fig.3d**) further confirmed a significant reduction in task accuracy following TTX treatment, validating that the observed behaviors were driven by active neural processing within the organoid. Together, these results demonstrate that Brainobot can perform multi-class decision-making tasks, highlighting the potential of organoid processors for task-specific robotic control.

**Figure 3.**
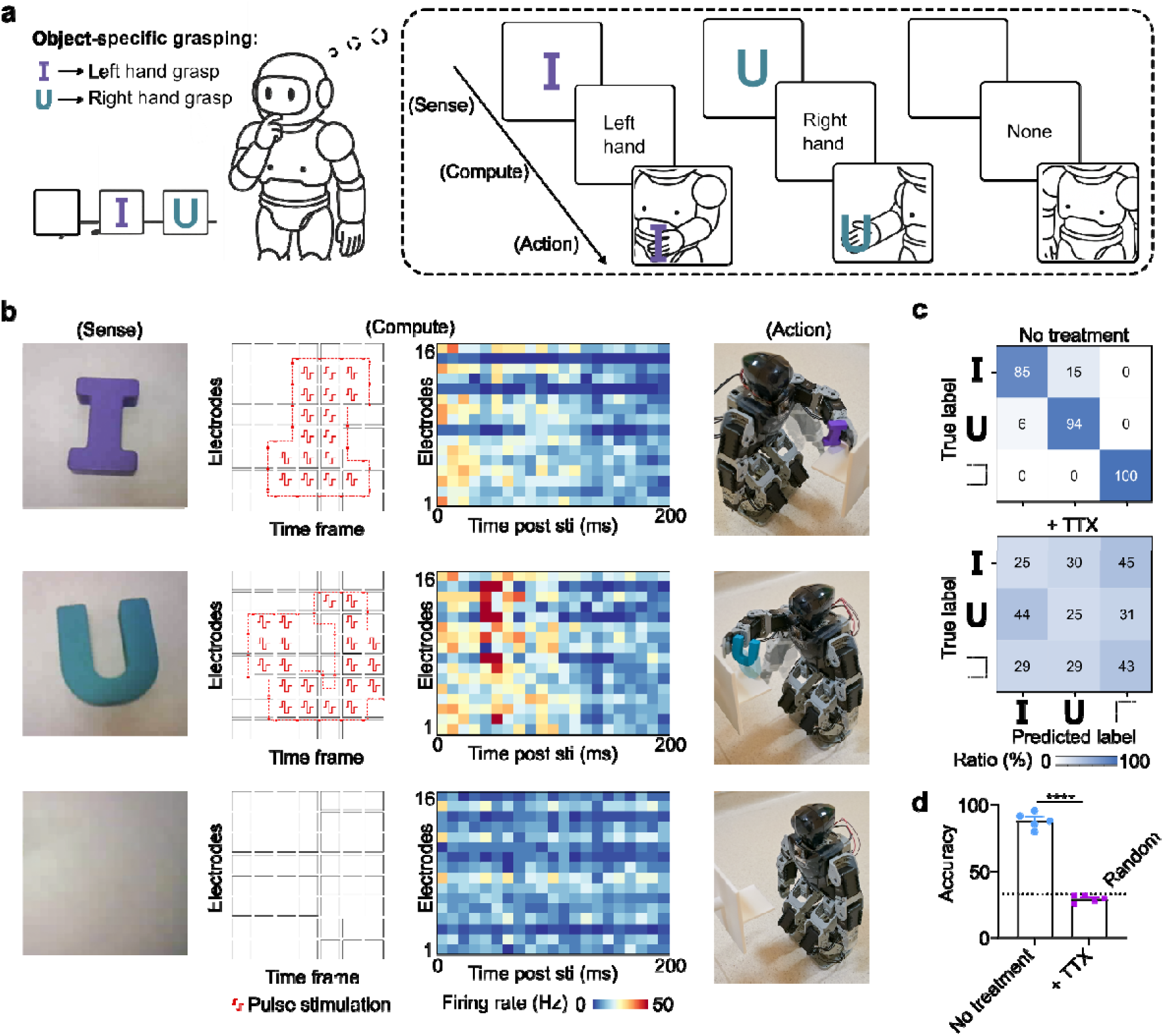
Grasping task. **a.** Design of the object-specific grasping task. The humanoid grasps the letter block “I” using its left hand, grasps the letter block “U” using its right hand, and performs no action when no letter block is presented. **b.** Conduction of the object-specific grasping task through a sense–compute–action loop. Representative images of the letter blocks (e.g., “I”, “U”, and blank) captured by the humanoid camera. Temporospatially encoded stimulation patterns corresponding to the three visual cues, together with representative raster plots showing the evoked neural activity in response to the three stimulation patterns. Representative images showing the humanoid grasping the letter block “I” with its left hand, grasping the letter block “U” with its right hand, or performing no action in response to the corresponding visual cues. **c.** Representative confusion matrices showing the performance of the organoid processor without and with TTX treatment. **d.** Quantification of organoid processor performance without or with TTX treatment (meann±ns.e.m., *no*=D5 organoids, from three independent experiments).

### Laser chasing task

We further extended the application of Brainobot to enable interaction with external environments and perform continuous, multi-step decision-making tasks. As a proof-of-concept example, we designed a laser-chasing task involving sequential motor actions (**Fig.4a**). In this task, Brainobot was programmed to pursue the location of a laser spotlight through a closed-loop sense–compute–action framework. Within this framework, the humanoid used onboard cameras and sensors to capture environmental images and distance information (Sense), processed this information using the organoid processor to generate high-level commands including turning left, turning right, or moving forward (Compute), and transmitted these commands to mid-/low-level controllers to drive the humanoid’s robotic legs for locomotion (Action) (**Fig.4b, Supplementary Fig.7, Supplementary Video.2**). Specifically, Brainobot was programmed to make decisions every 5 seconds at each step until the humanoid ultimately reached the laser spotlight target. During each step, the organoid processor identified the location of the spotlight by receiving encoded environmental image features from the PC through the MEA interface, generated distinct patterns of evoked neural activity corresponding to different spotlight locations, and output decoded control signals for locomotion movements of the humanoid. Through this multi-step decision-making process, the humanoid successfully approached the laser spotlight target under the guidance of neural activity generated by the organoid processor (**Fig.4c, Supplementary Fig.8, Supplementary Video.3**). The confusion matrix (**Fig.4d**) showed that the organoid processor achieved high accuracy in the laser-chasing task, whereas TTX treatment markedly impaired task performance (**Supplementary Video.4**). Quantitative analysis (**Fig.4e**) further demonstrated a significant reduction in task accuracy following TTX treatment, supporting that the observed behaviors were mediated by active neural processing within the organoid processor. Together, these results demonstrate that Brainobot can achieve closed-loop, real-time interaction with external environments.

**Figure 4.**
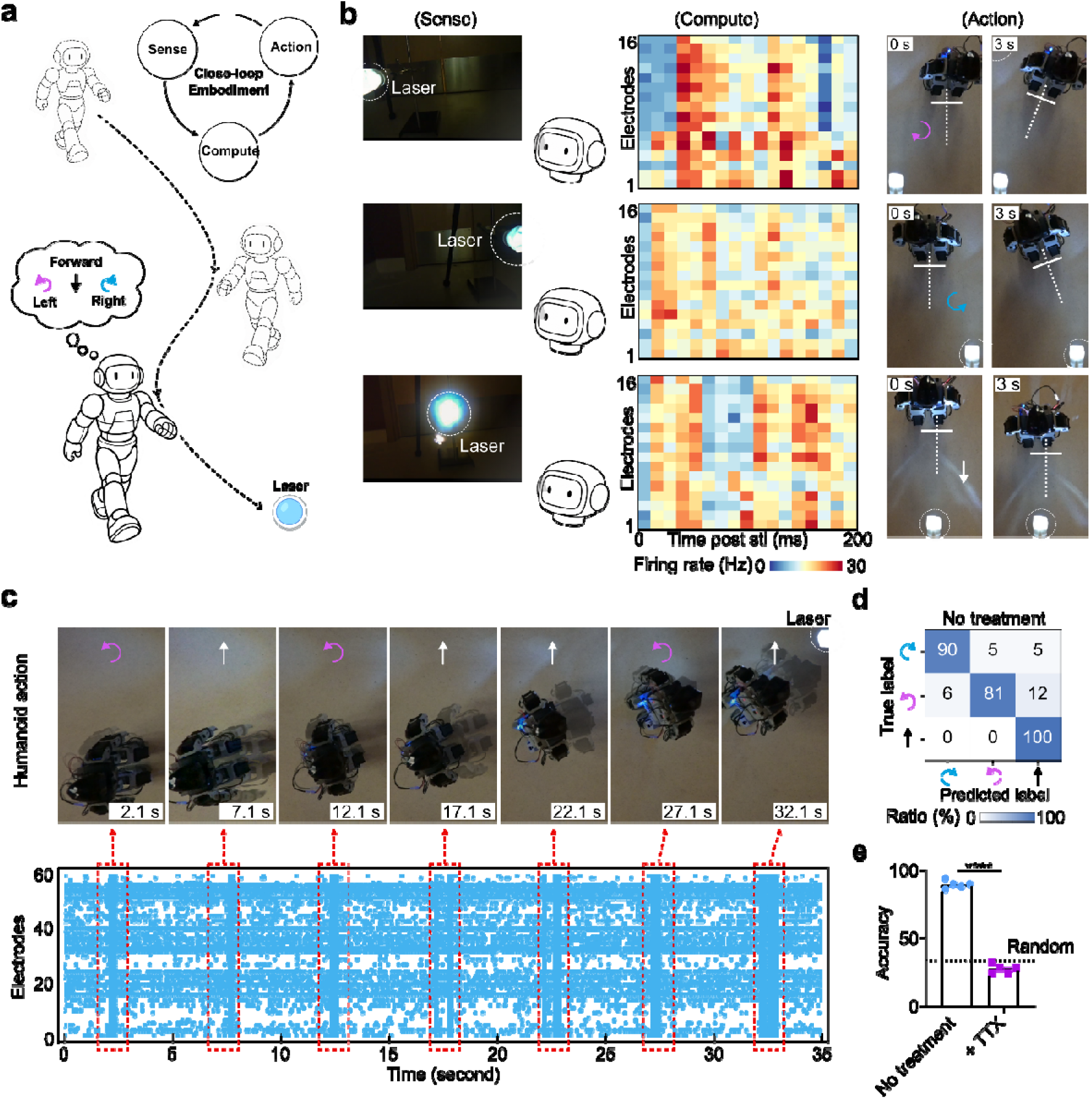
Laser chasing task. **a.** Design of the laser chasing task. The humanoid moves forward when the laser pointer is detected in front, turns right when the laser dot appears on the right, and turns left when the laser is on the left. **b.** Representative images of the laser pointer in different locations relative to the humanoid, together with representative raster plots showing the evoked neural activity in response to corresponding sensory inputs, and representative images of final humanoid motor actions. **c.** Overview of a complete laser chasing process. Time-lapse images showing the humanoid chasing the laser procedure (Top) and correlated neural activities (e.g., raster plots) of the organoid processor over the full task duration (Bottom). **d.** Representative confusion matrix describing the task performance. **e.** Quantification of organoid processor task performance without or with TTX treatment (meanL±Ls.e.m., *no*=D5 organoids, from three independent experiments).

### Cross-task adaptivity and high-efficiency learning

Building on these proof-of-concept task demonstrations, we further investigated unique features of Brainobot, including cross-task adaptivity and high-efficiency learning. To evaluate these properties, we designed three experimental groups under different conditions, with one training epoch performed every half day (**Fig.5a**), and subsequently assessed Brainobot performance accordingly (**Fig.5b, Supplementary Fig.9**). The representative learning curve of Brainobot (**Fig.5b**, Group 1) showed that task accuracy first increased from 72% to 92% during three sequential laser-chasing tasks (epochs 0, 1, and 2), decreased to 86% after switching to the grasping task (epoch 3), and then further increased to 92% during three sequential grasping tasks (epochs 3, 4, and 5), suggesting adaptive learning in Brainobot.

**Figure 5.**
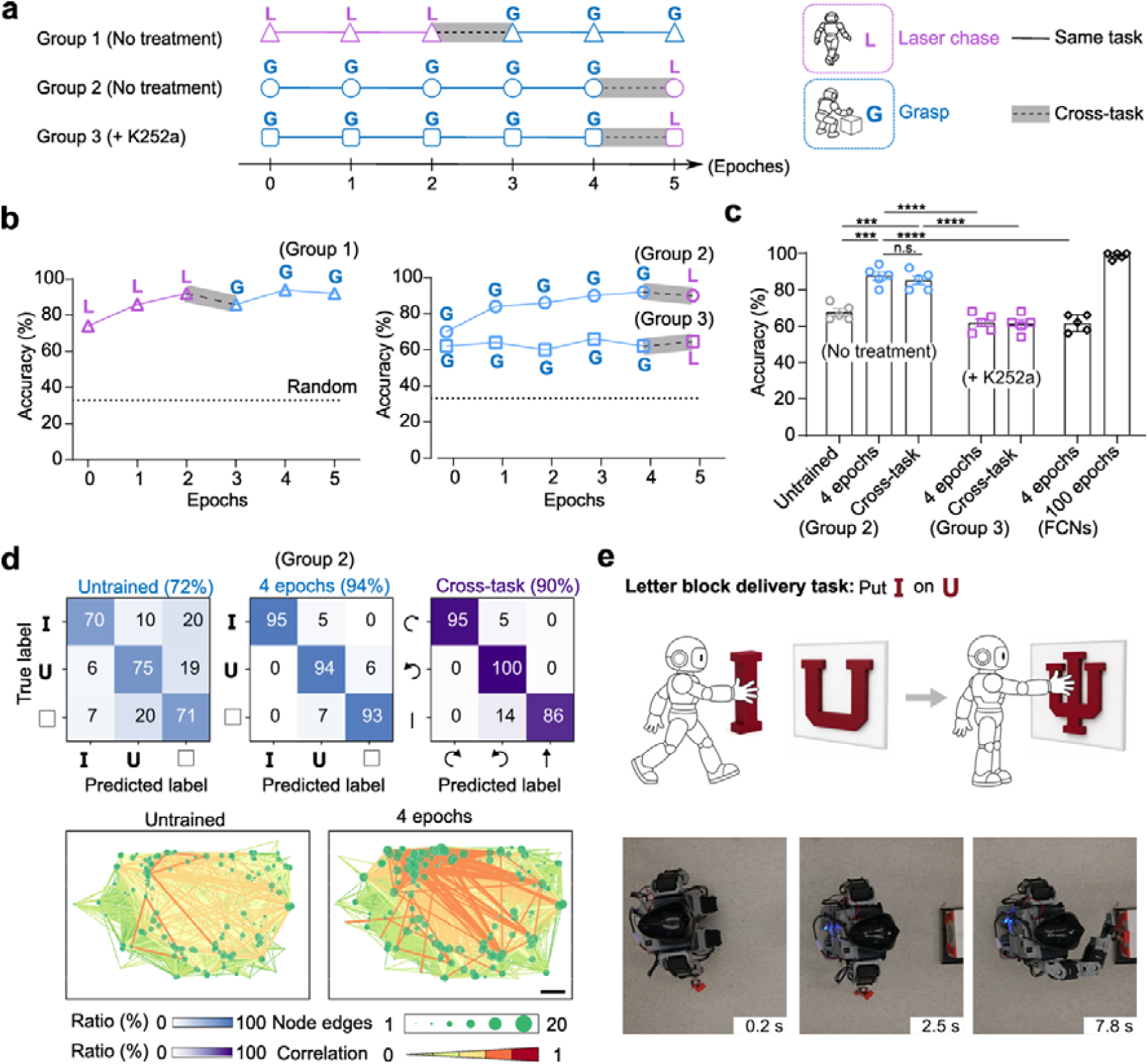
Cross-task adaptivity and high-efficiency learning. **a.** Schematic illustration of the experimental design for three training groups across different tasks (e.g., different combinations and sequences of laser-chasing and object-grasping tasks) and treatment conditions (e.g., with or without K252a treatment). **b.** Representative learning curves of the three training groups demonstrating cross-task adaptivity. **c.** Quantification of Brainobot performance under different stages and conditions (mean ± s.e.m., n = 5 organoids from three independent experiments), together with comparison to fully connected networks (FCNs), indicating high-efficiency learning. **d.** Representative confusion matrices showing grasping performance throughout the training process and laser-chasing performance after cross-task adaptivity (top), together with the corresponding functional connectivity maps during training (bottom). **e.** Design and demonstration of a sequential multi-stage task: “Put I on U” to form the “IU” logo.

Interestingly, the trained Brainobot following three sequential laser chasing tasks (Group 1, epoch 3, 86%) exhibited significantly higher accuracy than the untrained Brainobot (Group 2, epoch 0, 70%) when performing the same grasping task, indicating cross-task adaptivity. Moreover, an additional representative learning curve of Brainobot (**Fig.5b**, Group 2), together with the corresponding quantitative analysis (**Fig.5c**, Group 2), further confirmed cross-task adaptivity and suggested that this capability was largely independent of the training sequence. Furthermore, compared with a representative machine learning algorithm, a fully connected network (FCN), Brainobot achieved superior performance under the same training condition of four epochs in the grasping task (**Fig.5c**, Group 2 vs FCNs), suggesting high-efficiency learning of Brainobot.

The pronounced learning behavior and cross-task adaptivity observed in Brainobot may be associated with activity-dependent synaptic plasticity within the brain organoid neural networks. To investigate this possibility, we performed the same training procedure in the absence (Group 2) or presence (Group 3) of K252a treatment, a kinase inhibitor commonly used to suppress activity-dependent synaptic plasticity, and analyzed the functional connectivity of the organoid processor. Representative confusion matrix results (**Fig.5d**, top) showed that the accuracy of Brainobot (Group 2) first increased from 72% to 94% during five sequential grasping tasks (epochs 0, 1, 2, 3, and 4), and then decreased to 90% after switching to the laser-chasing task (epoch 5). Correspondingly, apparent changes in functional connectivity were observed throughout the training process (**Fig.5d**, bottom). In contrast, the accuracy of Brainobot with K252a treatment (**Fig.5b**, Group 3) remained between 62% and 64%, while functional connectivity changes were minimal due to inhibition of synaptic plasticity (**Supplementary Fig.9b**).

We further investigated the capability of Brainobot p to conduct sequential multi-stage tasks using a letter-block delivery task, “Put I on U,” to form the “IU” logo (**Fig.5e, Supplementary Video.5**). In this composite task, the humanoid was programmed to carry the letter block “I” in its right hand, locate the letter block “U” at a target position, and subsequently place the block “I” onto “U” to form the “IU” logo. Brainobot successfully completed this multi-stage task, resulting in the formation of the complete “IU” logo through coordinated sequential actions. These results demonstrate the capability of Brainobot to integrate and execute complex sequential multi-stage tasks.

## Discussion

In this study, we present Brainobot, a biohybrid robotic platform that integrates living cortical organoids as adaptive controllers for autonomous decision-making. By leveraging reservoir computation of organoid neural networks, Brainobot enables real-time sensory processing and action control through a closed-loop embodiment between organoid activity and humanoid. This system supports complex robotic behaviors in a physical world, including grasping and laser chasing, meanwhile also exhibits emergent features of biological intelligence including adaptivity, generalization, and self-organization stemming from experience-dependent network dynamics(*21, 46*). These results establish a framework for constructing biohybrid robotic systems that extend beyond current silicon-based computing hardware systems toward biologically embodied intelligence.

Brainobot introduces a new platform for high efficiency, adaptive decision-making by leveraging organoid intelligence for closed-loop physical embodiment, holding promising potential for the applications in autonomous robotics, biorobots, and synthetic intelligence systems. Beyond its engineering significance and robotic advances, it also provides a unique platform to explore biological intelligence by mapping neural network function to intelligent behaviors, which may offer new insights for understanding and treating neurological disorders using patient-specific organoids. Nonetheless, Brainobot faces several important limitations. First, current brain organoids face challenges including heterogeneity, hypoxia/necrosis, and issues on reproducibility, scalability, and organoid/MEA interface,(*47–49*),(*50*) limiting the system performance. Engineering advances in microphysiological systems, AI-enabled microreactors, and soft bioelectrodes may address these challenges in the future(*51–54*). Second, despite the unique strength of Brainobot, the supporting infrastructure including incubators, MEA systems, and external computation remains bulky and energy-intensive, requiring engineering efforts for system miniaturization and optimization. Furthermore, efficient encoding and decoding of real-time information remain difficult, and advanced computational frameworks will be greatly needed to manage, analyze, and interpret the large volumes of data generated by organoid-based systems. Finally, ethical considerations will require careful attention as biohybrid systems become increasingly intelligent (*55*). Ethically appropriate use cases and clear public communication will have to be considered as this technology advances.

## Methods

### Human stem cell culture

We obtained the H9 hESC (human embryonic stem cell) line from WiCell Institute. We followed the guidelines from the WiCell Institute and the Indiana University Biosafety Committee. We cultured the ESCs on 6-well plates with GFR (growth factor-reduced) Matrigel (Corning). We maintained the ESCs in a humidified incubator at 5% CO_2_ and 37°C with medium (mTeSR plus, Stem cell technology) changed every 2 days. We used ReLeSR (Stemcell Technologies) for passaging the ESCs when they reach 70-80% confluence.

### Generation of cortical organoids

The cortical organoids were generated using our previously established protocol(*40, 56, 57*), which was adapted from published protocols(*36, 37*). In brief, we generated embryoid bodies (EBs) by introducing 4 million stem cells into one well of 6-well ultra-low attachment plates and shaking at 90 RPM orbital shaking. The EBs were cultured in the mTeSR Plus medium (STEMCELL Technologies) supplemented with 10 μM SB431542 (Stemgent), 1 μM Dorsomorphin (R&D Systems), and 10 μM Y-27632 (Only for 24 hours, SelleckChem). After three days, the developing structures were moved to cortical organoid maintenance medium (Neurobasal (Life Technologies) with GlutaMAX, 1% Gem21 NeuroPlex, 1% N2 NeuroPlex (both from Gemini Bio-Products), 1% non-essential amino acids (NEAA), 1% penicillin-streptomycin (P/S)), supplied with 10 μM SB431542 and 1 μM Dorsomorphin. On Day 10, cultures were switched to cortical organoid maintenance medium supplemented with 20 ng/mL FGF2 (Life Technologies) for 7 days. This was followed by another 7 days in cortical organoid maintenance medium supplemented with both 20 ng/mL FGF2 and 20 ng/mL EGF (PeproTech). Subsequently, the organoids were cultured in cortical organoid maintenance medium supplied with 10 ng/mL each of BDNF, GDNF, and NT-3 (PeproTech), along with 200 μM L-ascorbic acid and 100 μM db-cAMP (Sigma-Aldrich) to support neural maturation. After 7 days, organoids were maintained in cortical organoid maintenance medium without any supplements.

### Attach to Maxone MEA

Two-month-old organoids were used for MaxOne MEA attachments. Before attachment, MaxOne chips were first treated with 0.07% polyethylenimine (PEI) in DI water for 1 hour, followed by incubation with 10 μg/mL human laminin-521 in organoid maintenance medium for 30 minutes to promote cellular attachment. The plated organoids were initially maintained in cortical organoid maintenance medium. On 3 months, the medium was switched to BrainPhys medium (STEMCELL Technologies) supplemented with 2% NeuroCult™ SM1, 1% penicillin-streptomycin (P/S), 10 ng/mL BDNF, GDNF, and NT3. The attached organoids were maintained in standard humidified incubator at 5% CO_2_ and 37°C.

### Neural activity measurements

The neural activity measurements were conducted using MaxOne MEA with the MaxLab Live Scope <u>V20.1</u>. The MaxOne system has 26400 electrodes with 1024 electrodes recorded simultaneously. The system was set to 512x gain with a 300 Hz high pass filter, and 5.5x std threshold for spike detection, as recommended by the manufacturer. The organoids were recorded 2 to 24 hours after medium change. To locate the active electrodes with neural activities from the cortical organoids, we first conducted an activity scan using the Checkerboard configuration in the MaxLab Live Scope. After the activity scan, an activity map of the organoid that showed the active area of spontaneous spiking activities could be obtained. Based on the activity map, we selected the top active electrodes (mean firing rate > 0.2Hz, maximum 1024 electrodes) for spontaneous network activity recordings and organoid computing.

### Spike sorting

As the electrode of the MaxOne chip is 12.5 um and smaller than the length of neural processes, activity from one neuron could be detected from multiple electrodes. To obtain single-unit spontaneous activity, we performed spike sorting following our previously established protocol. In brief, the raw recording was first filtered using a 300-6000Hz band pass filter and then spike sorted using SpikeInterface framework and Kilosort2 software with the default parameters. We concatenated the recordings from the same chip with the same electrode configuration on the same day for spike sorting.

### Functional connectivity

We performed 5-minute spontaneous activity recordings for the functional connectivity analysis. The raw recordings were first spike sorted. The functional connectivity of human cortical organoids was calculated from the sorted spontaneous activity recording using an STTC (Spike time tiling coefficient) method. We calculated the STTC in a time window of 30ms and used a threshold of 0.35 to rule out most random connectivity. Custom MATLAB code adapted from a previously published work was used to calculate the STTC correlation, and a custom Python script was used to plot and visualize the functional connectivity map.

### Electrical stimulation

We employed the Python API of MaxLab Live to deliver spatiotemporal electrical stimulations to cortical organoids. Based on the activity scan, we selected the electrodes with the largest spike amplitude and minimal distances between electrodes larger than 100 μm for electrical stimulation. Based on our previous optimization, we employed bipolar pulse stimulation with pulse time = 500 μs, pulse interval = 100 ms, and pulse voltage = 100 ∼ 500 mV. The detailed stimulation parameters and encoding for different tasks can be found below.

### Hardware integration of Brainobot

The Brainobot system integrates organoid electrophysiology, visual sensing, and motion control into a fully wireless closed-loop platform (**Fig.S3a**). First, a camera module (ESP32-CAM) provided visual perception by continuously streaming image data to the host PC, where object detection and positional tracking were performed in real time. These visual features were then converted into patterned electrical stimuli delivered to the organoid via a multi-electrode array (MaxOne), which simultaneously recorded neural responses. The recorded electrophysiological activity was processed on the PC to generate discrete decision outputs representing behavioral intentions. Finally, the decoded commands were transmitted through a Bluetooth transceiver pair (BT-410) to the robot’s main controller (CM-530), which executed corresponding motion primitives from a predefined library using Dynamixel AX-12A actuators. The robot was powered by an 11 V Li-Po battery, while a DMS-80 infrared sensor provided distance feedback for basic obstacle avoidance. Together, these modules established a seamless perception–computation–action loop between the living neural network and the robotic embodiment.

### Software integration of Brainobot

The Brainobot control framework was organized into three functional layers: data acquisition, data computation, and robot control (**Fig.S3b**). In the data acquisition layer, visual information was obtained through a camera module and processed on the host computer using custom Python scripts. The incoming frames were first subjected to color segmentation using hue–saturation–value (HSV) filtering and blue-ratio enhancement to isolate the target object. When color segmentation failed under low-light conditions, the pipeline automatically switched to a grayscale thresholding fallback. Morphological filtering was applied to remove noise and refine object contours, followed by cropping, uniform scaling, and center alignment. The extracted spatial information was then mapped into a stimulation pattern (“Stim Mapper”) that defined the organoid input channels on the MEA interface. In the data computation layer, the organoid received patterned electrical stimuli and generated spontaneous neural responses. Electrophysiological signals were continuously recorded, filtered, and processed in Python for spike detection and rate estimation. The resulting spike trains were converted into a real-time state vector, which was analyzed by a readout layer to classify the ongoing neural dynamics into discrete decision outputs (e.g., “no object” or “move toward object”). In the robot control layer, the decoded action was transmitted to the robot’s CM-530 controller for execution. The controller operated a hierarchical planner that translated high-level decisions into corresponding motion primitives drawn from an onboard motion library. Mid-level routines handled sequencing and servo dispatch, while low-level control executed precise actuator commands via Dynamixel motors and returned a task-completion flag. A brief delay was introduced before resuming the next sensing cycle, forming a continuous perception–computation–action loop.

### Encoding for the grasp task

The raw images captured by the camera on the humanoid were first cropped and transmitted into 7 x 8 binary images using a threshold of 128. Each 7×8 binary image was converted into 8 pulse streams, with each row represented by a pulse stream. Then, the pulse streams were applied to the 6 stimulation electrodes (selected based on the description in the ‘Electrical Stimulation’ session). The pulse streams were stimulated to the cortical organoids at 100-ms intervals, and each 7 pulse streams corresponding to each training image was stimulated to the organoids at 5-second intervals. The evoked activity after the stimulation of each pulse train was recorded, and the number of spikes in 100-ms time bins after the stimulation was obtained as the reservoir state for further decoding to get the classification output.

### Implementation of the grasp task on Brainobot

The grasp task was designed to demonstrate closed-loop perception–computation–action control mediated by organoid computation. The ESP32-CAM module first captured a static image of the workspace, which was processed through the Python-based visual pipeline described above. The image underwent color segmentation, morphological filtering, and feature extraction to isolate the target region containing an alphabetic cue. The resulting spatial pattern was transformed into an electrical stimulation map and delivered to the organoid via the MEA interface. The organoid’s electrophysiological responses were recorded, filtered, and analyzed to classify the stimulus into one of three decision states: detection of “I”, detection of “U”, or no detection. Each state was mapped to a corresponding motor command: detection of “I” triggered a left-hand grasp, detection of “U” triggered a right-hand grasp, and the absence of either symbol resulted in a standby posture without actuator movement. The decoded motor command was transmitted via Bluetooth to the controller, which executed the appropriate motion primitive using actuators. Real-time joint feedback and distance sensing ensured stable grasp execution and object contact detection.

### Encoding for the laser chasing task

The raw images captured by the camera were first cropped and transformed into 6 x 12 binary images using a threshold of 128. Each 6×12 binary image was reshaped into a 12 x 6 binary image and then converted into 12 pulse streams, with each row represented by a pulse stream. The pulse streams were stimulated to the cortical organoids at 100-ms intervals, and each 12 pulse streams corresponding to each training sensory input of the laser position was stimulated to the organoids at 5-second intervals.

### Implementation of the laser chasing task on Brainobot

The laser chasing task was implemented to demonstrate dynamic sensorimotor control and real-time adaptation in the organoid-driven loop. A laser dot projected within the robot’s visual field was captured by the ESP32-CAM module and processed through the Python-based image analysis pipeline. The centroid of the detected laser spot was calculated relative to the image center and transformed into a spatial stimulation pattern, which was delivered to the organoid through the MEA interface. The organoid’s neural responses were analyzed online to classify the stimulus into one of three decision states: left, center, or right. When the laser spot appeared on the left side, the organoid output triggered a rightward rotation to realign the robot toward the target; when the spot appeared on the right, a leftward rotation was executed. If the laser was centered, the robot initiated a forward movement to pursue the target. A distance feedback loop was integrated through the DMS-80 infrared sensor mounted on the robot’s torso. Once the measured distance between the robot and the target fell below a preset threshold, a stop command was automatically triggered to terminate movement and prevent collision. The classified motor command was transmitted to the CM-530 controller via Bluetooth (BT-410) and executed using Dynamixel AX-12A actuators.

### Read-out function

We decoded the evoked activities using the read-out function following our previously established protocol. In brief, evoked spiking activities from the top active electrodes were converted into a response matrix by binning the spikes within defined time windows. In this response matrix, the y-axis represents electrode indices, the x-axis corresponds to time bins, and each data point denotes the spike count at an electrode and time bin. This matrix was then flattened into a one-dimensional vector x, which served as the reservoir state and the input for the downstream logistic regression decoding algorithm, a simple function that applies a sigmoid function to the linear combination of the input vector **x** to obtain categorical outputs **y**:

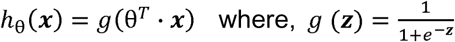

Model training was conducted using the Scikit-learn library in Python, and performance was evaluated using 5-fold cross-validation on a single epoch of training data.

### Training of Brainobot

The training of Brainobot was achieved by repeatedly electrical stimulation of the same input dataset to induce activity-dependent neural plasticity. For each training epoch, we gave electrical stimulation patterns corresponding to the entire dataset to the organoid bio-controller, and the ONNs reshaped the plasticity and reorganized the connectivity in response to the electrical stimulation. Following each stimulation cycle, we trained the readout function to adapt to the updated network dynamics, resulting in improved system performance.

### Staining

Organoids were rinsed in 1× PBS (Gibco)for 3 times, fixed in 4% paraformaldehyde (PFA; Thermo Scientific) overnight at 4□°C, and then transferred in 30% sucrose overnight at 4□°C. The fixed tissues were then snap-frozen in cryomolds (Sakura Finetek) with O.C.T. (Fisher Scientific) in dry ice. Sections (30Dpm thick) were obtained using a Leica cryostat and mounted onto charged slides. For antigen retrieval, sections were incubated in citrate buffer at 96°C for 20 minutes, washed, and blocked in PBS with 0.3% Triton X-100 and 5% normal goat serum for 1 hour. Primary antibodies were applied overnight at 4□°C in a humidified chamber. After PBS washes, secondary antibodies were incubated at room temperature for 2 hours. Nuclei were counterstained with DAPI and mounted using ProLong Gold Antifade medium (Invitrogen).

### Statistics

All analyses were performed using GraphPad Prism 8. Comparisons between two groups were made using Student’s *T*-test. Statistical significance was indicated as follows: p□<□0.05 (*), p□<□o.oi (**), p□<□0.005 (***), and p□<□o.ooi (****).

## Data Availability

All data are available in the main text or the supplementary figures.

## Acknowledgments

F.G. wants to acknowledge with the support from the National Science Foundation Award (EFRI2422149) and the National Institute of Health Awards (U54AG090792, R01GM160423).

## Author Contributions Statement

F.G. and Ho.C. conceived the study and designed experiments. Ho.C., C.T., Y.X., and Y. Y. performed the experiment with help from Z. H., Hu. C, J. W., and Z. A. H.C., C.T., Y.X., J.M., J.F., J.T., M.G., I.H., K.M., L.L., and F. G. analyzed the data. F.G. and Ho.C. wrote the manuscript. All authors read and provided feedback on the manuscript.

