## Supplementary material for "Brain organoid computing for robotic decision-making": SI

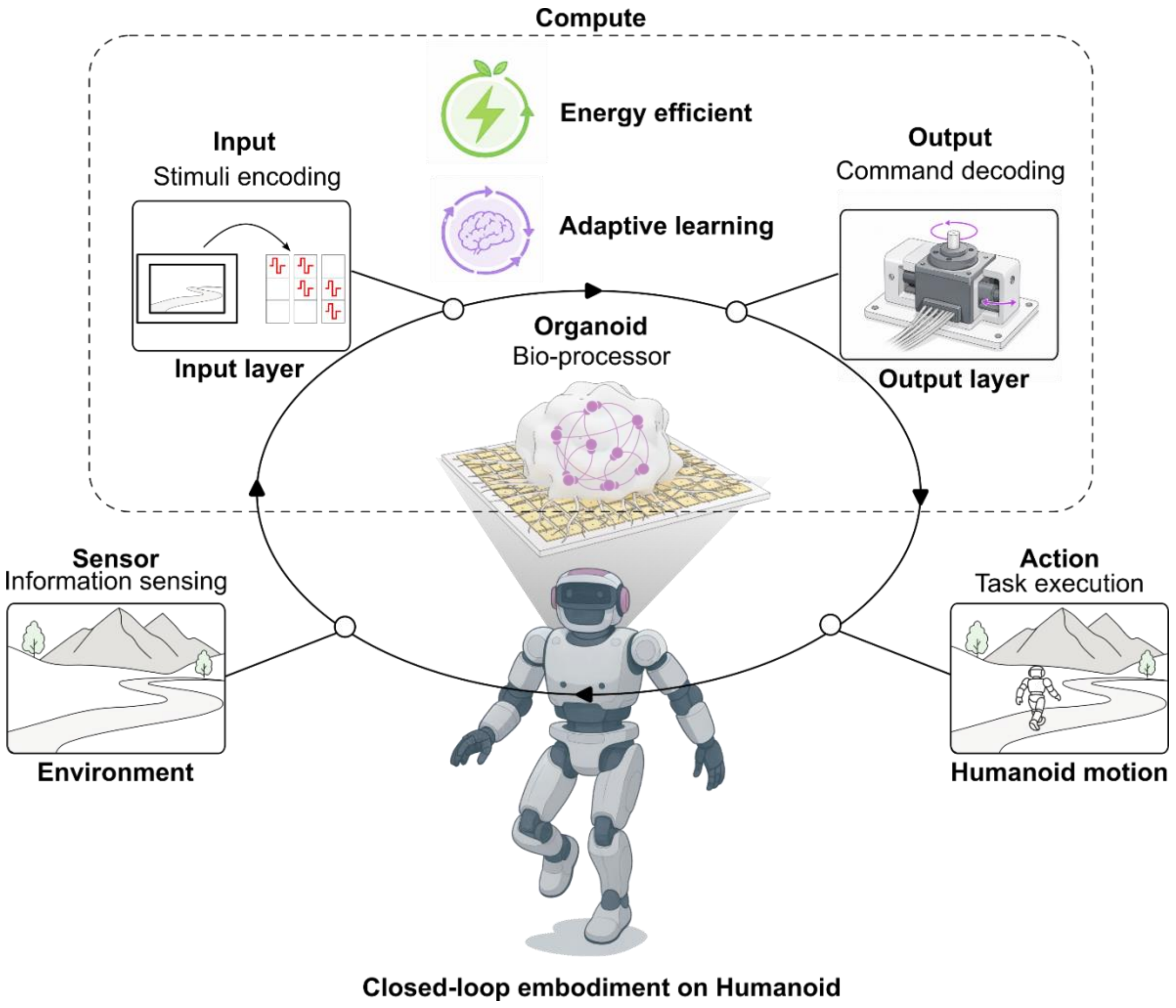


**Supplementary Fig. 1. Closed-loop sense-compute-action framework.** The brain organoid processor is integrated into the "Sense-Compute-Action" loop, enabling the humanoid to interact with the external environment.

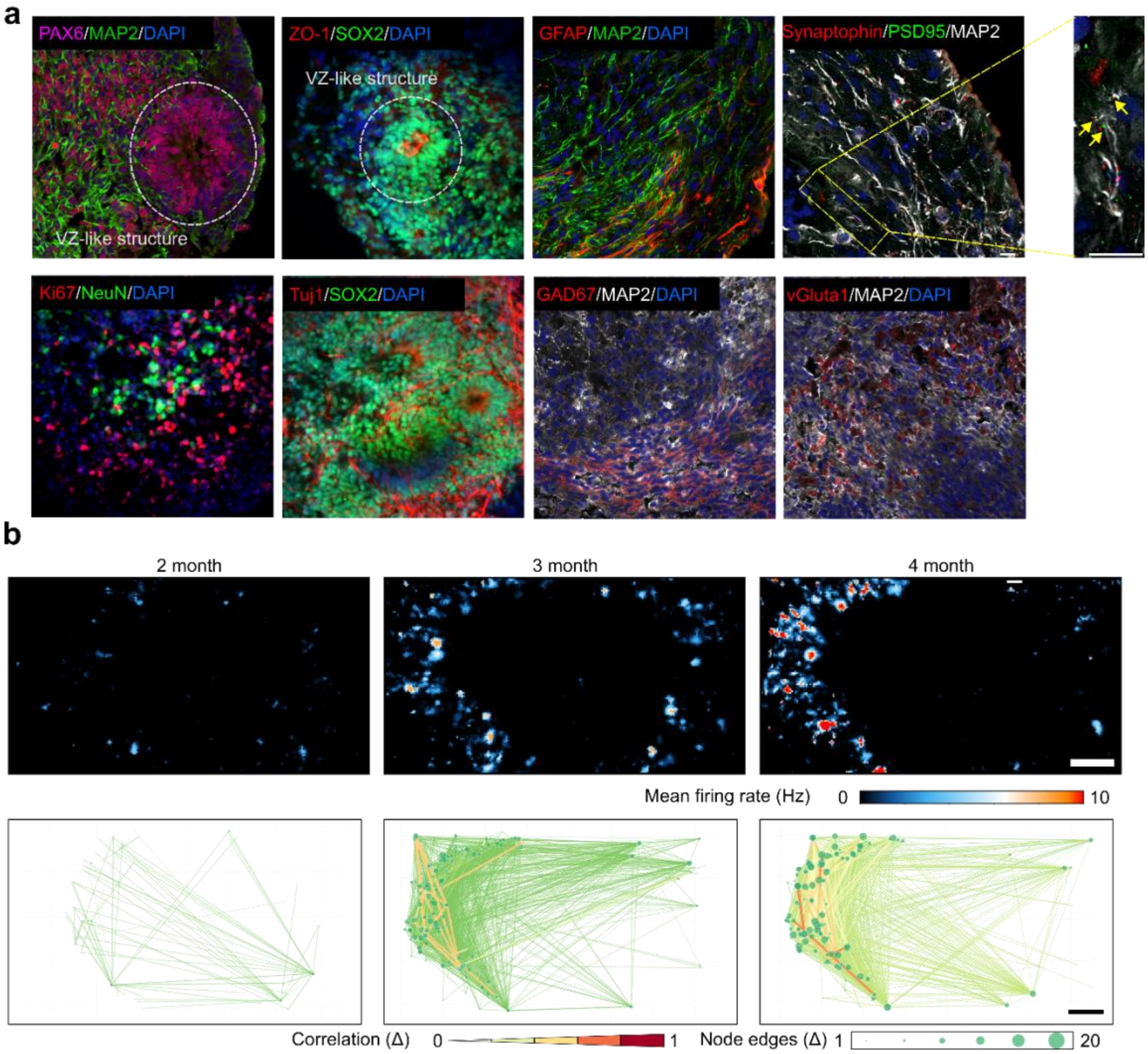


**Supplementary Fig. 2. Characterization of human cortical organoids.** (**a**) Representative immunofluorescent images showing the successful generation of human cortical organoids, scale bar = 20 μm. (**b**) Representative activity heatmaps and functional connectivity plots showing the functional maturation of ONNs, scale bar = 200 μm.

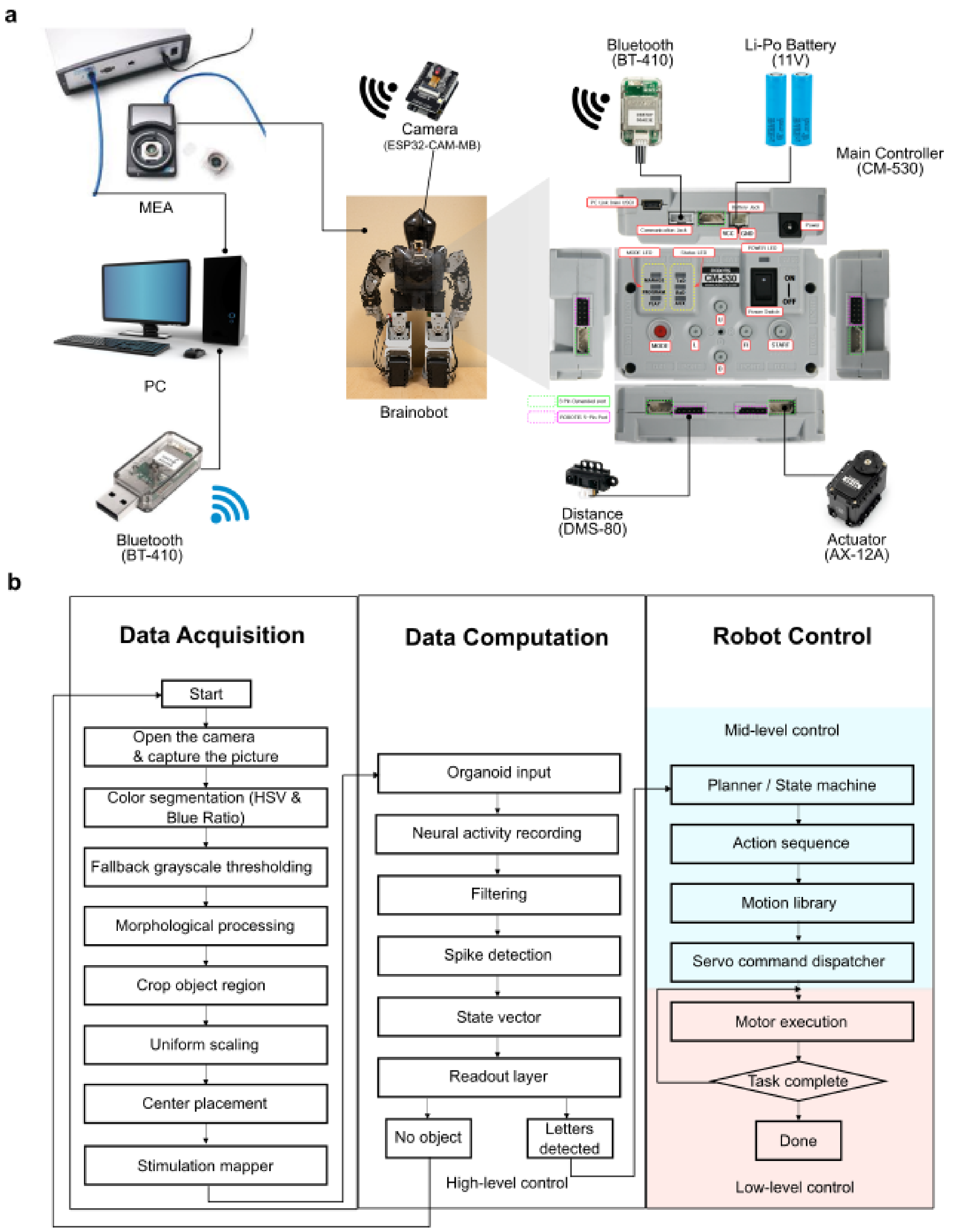


**Supplementary Fig. 3. System setup and control architecture of Brainobot.** Hardware architecture (**a**) handle signal acquisition and wireless communication, while software layers (**b**) implement data processing, organoid computation, and hierarchical robot control to realize a continuous perception–action loop.

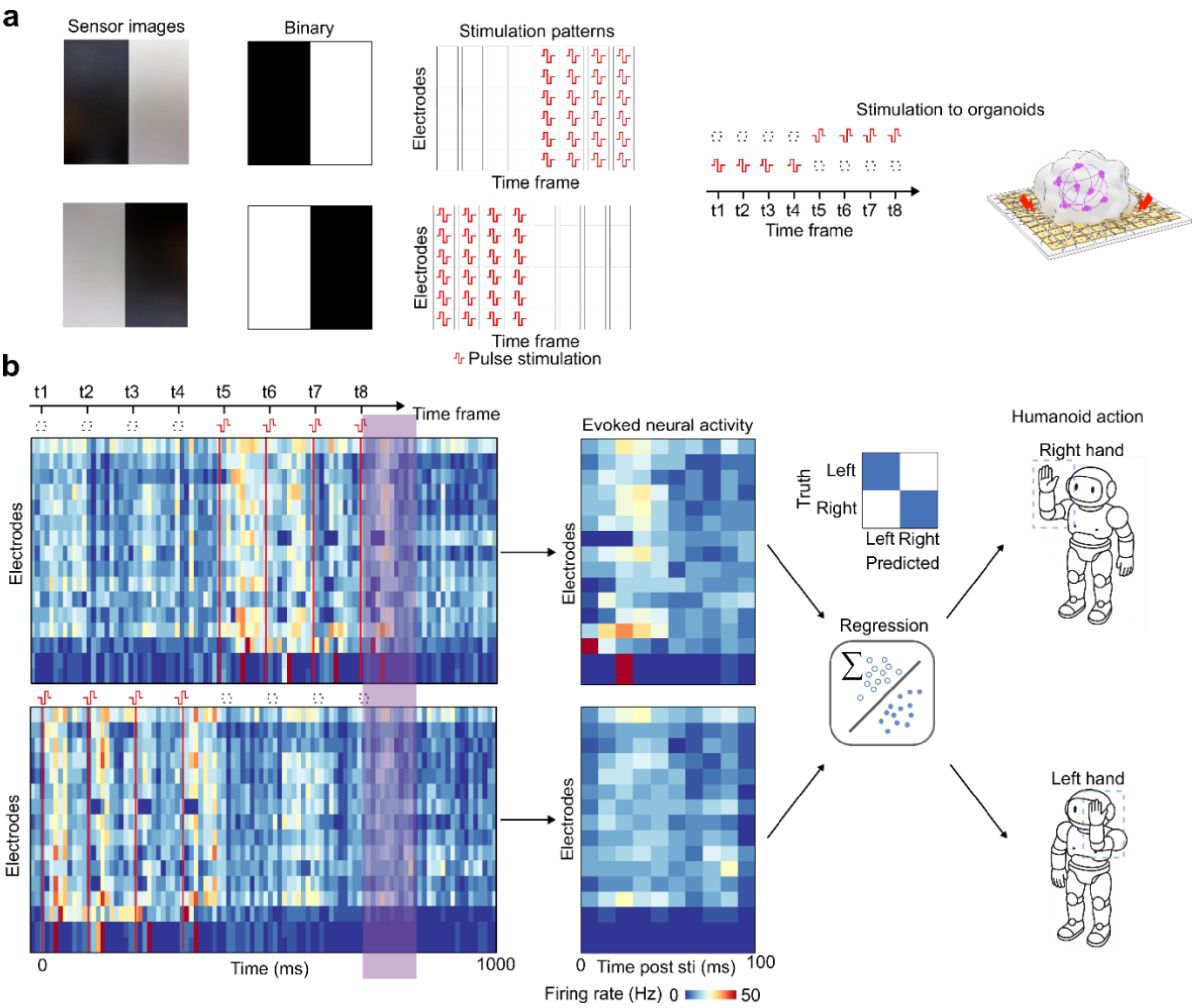


**Supplementary Fig. 4. Workflow of the cue-guided hand-raising task.** (**a**) Preprocessing of camera images into the temporal-spatial stimulation patterns for the organoid processor. (**b**) Output workflow of evoked activity from the organoid processor to the motion command of Brainobot.

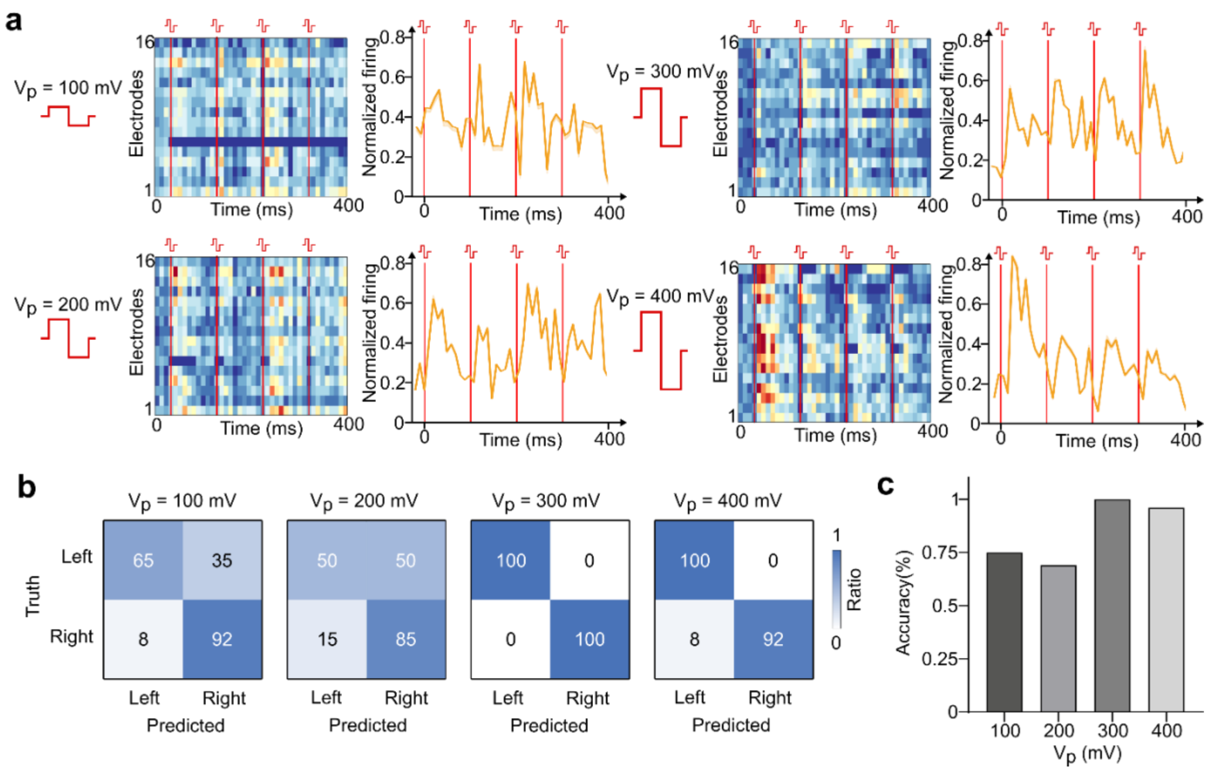


**Supplementary Fig. 5. Optimization of stimulation parameters.** (**a**) Raster plots and post-stimulation histogram showing the evoked activity responses of different voltages (Vp) by a stimulation sequence with four pulses. (**b**) Representative confusion matrix of the classification performance with four different stimulation voltages. (**c**) Bar plots showing the classification performance with four different stimulation voltages.

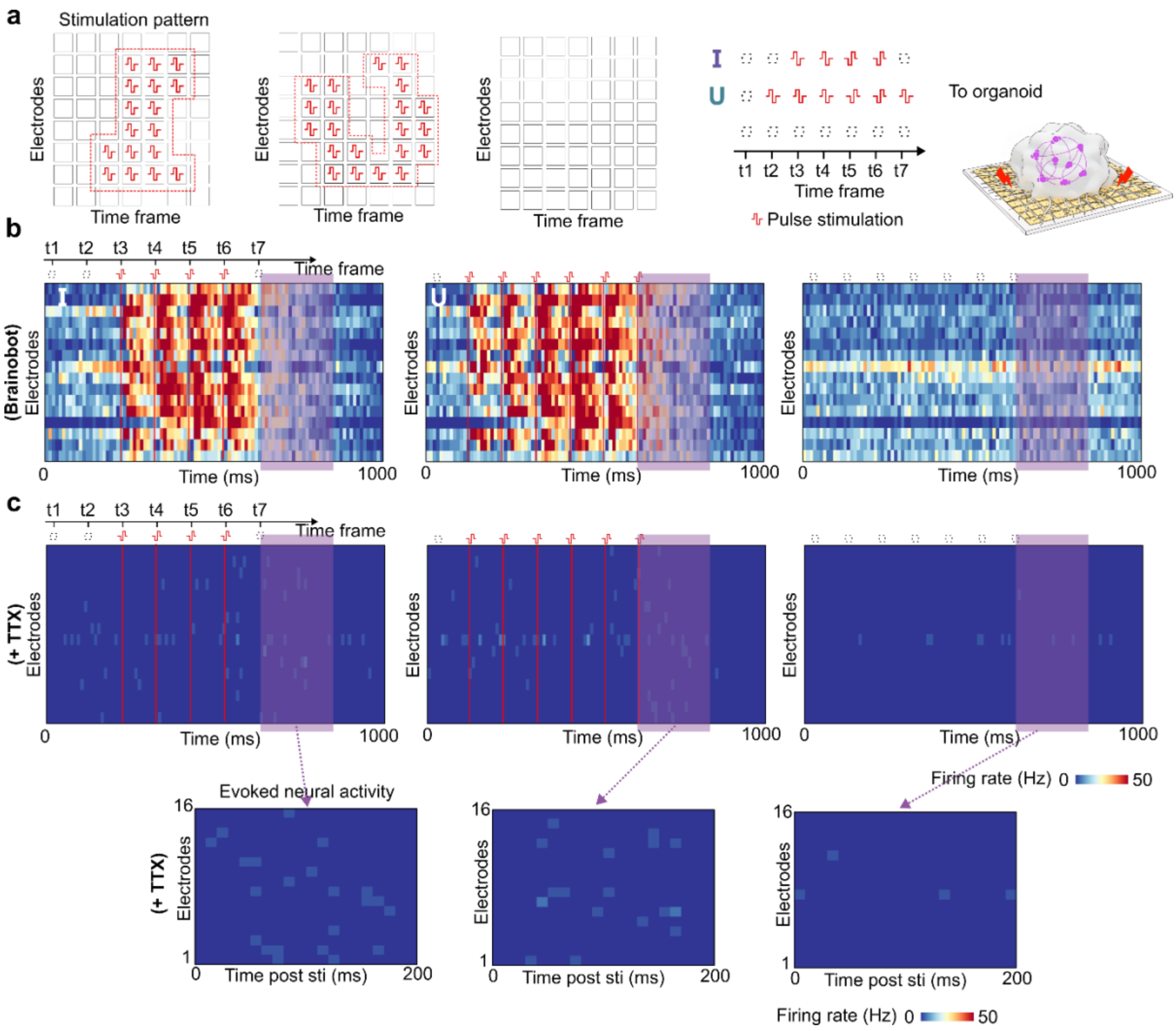


**Supplementary Fig. 6. Additional results related to the object-specific grasping tasks.** (**a**) Input workflow of sensory information to the organoid processor. (**b**) Evoked activity of the organoid processor in response to the 3 different stimulation sequences. (**c**) Evoked activity of the organoid bio-processor with TTX treatment in response to the 3 different stimulation sequences. (**d**) Evoked neural activity of the organoid processor with TTX treatment in response to the 3 different stimulation sequences.

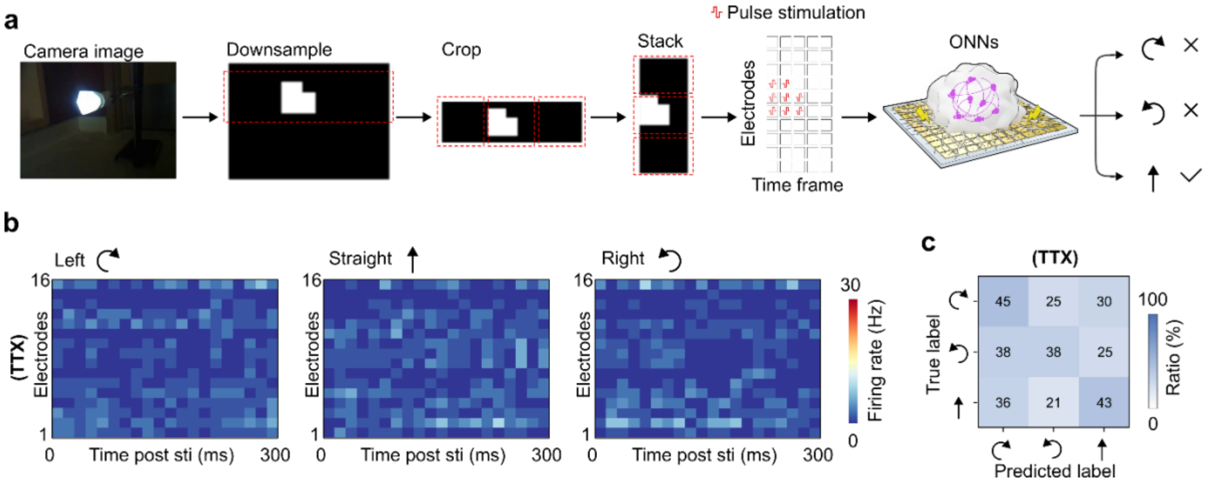


**Supplementary Fig. 7. Additional results related to the laser chasing task.** (**a**) Input workflow of sensory information to the organoid processor. (**b**) Evoked activity of the organoid processor treated with TTX in different input conditions. (**c**) Confusion matrix showing the performance of the organoid processor treated with TTX.

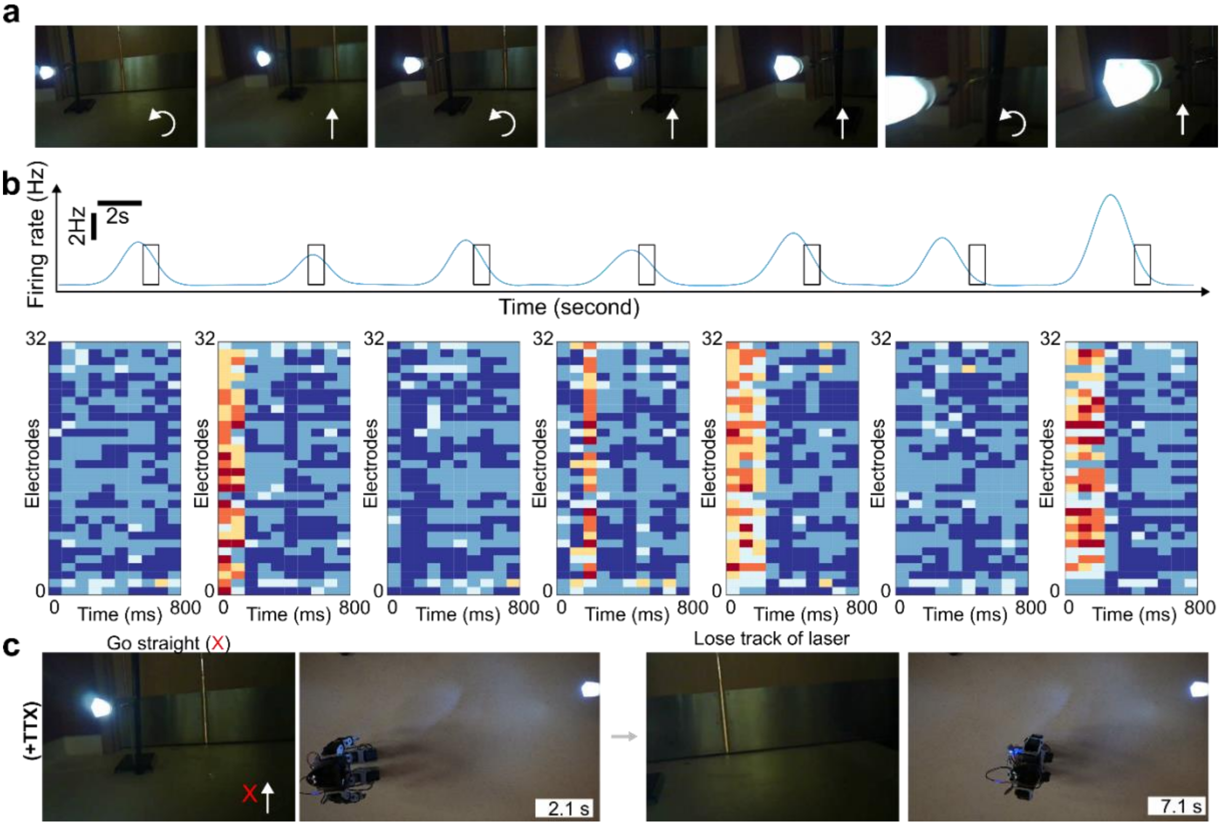


**Supplementary Fig. 8. Additional results related to the laser chasing task.** (**a**) Representative images showing the laser pointer over a complete laser chasing process captured by the camera sensor. (**b**) Histogram and raster plots showing the evoked activity of the organoid processor over a complete laser chasing process, corresponding to **a.** (**c**) Images showing the organoid processor treated with TTX made a wrong decision of "Go straight" when the laser was located on the left side of the humanoid.

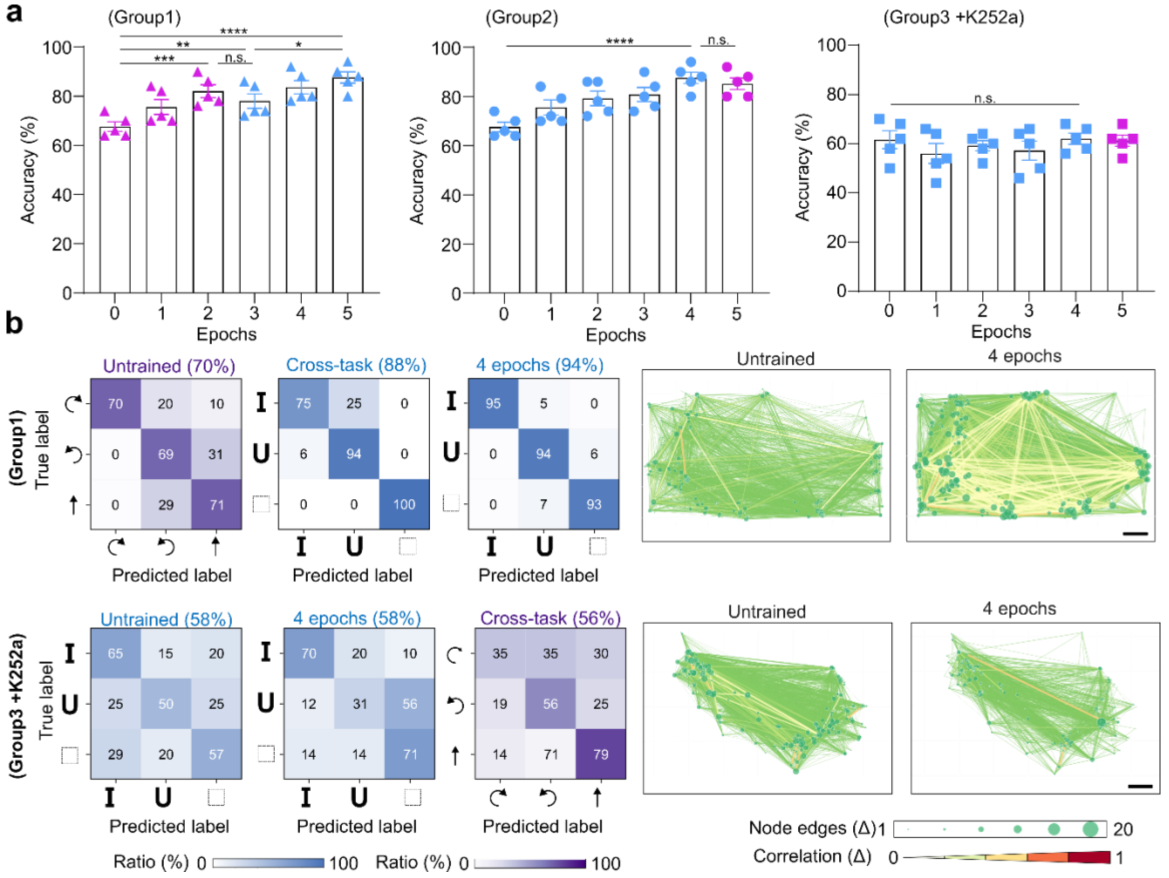


**Supplementary Fig. 9. Additional results related to adaptive learning.** (**a**) Performance during the training process of the three groups. (mean ± s.e.m., $n$ = 5 organoids, from three independent experiments). (**b**) Representative confusion matrix showing the grasp performance along the training process and the laser chasing performance after cross-task adaptivity of Brainobot with different training procedures (Group 1). Right, functional connectivity maps over the training process. Representative confusion matrix showing the grasp performance along the training process and the laser chasing performance after cross-task adaptivity of Brainobot treated with K252a (Group 3). Right, functional connectivity maps over the training process.
